# RefCell: Multi-dimensional analysis of image-based high-throughput screens based on ‘typical cells’

**DOI:** 10.1101/325415

**Authors:** Yang Shen, Nard Kubben, Julián Candia, Alexandre V. Morozov, Tom Misteli, Wolfgang Losert

## Abstract

**Background:** Image-based high-throughput screening (HTS) reveals a high level of heterogeneity in single cells and multiple cellular states may be observed within a single population. Cutting-edge high-dimensional analysis methods are successful in characterizing cellular heterogeneity, but they suffer from the “curse of dimensionality” and non-standardized outputs.

**Results:** Here we introduce RefCell, a multi-dimensional analysis pipeline for image-based HTS that reproducibly captures cells with typical combinations of features in reference states, and uses these “typical cells” as a reference for classification and weighting of metrics. RefCell quantitatively assesses the heterogeneous deviations from typical behavior for each analyzed perturbation or sample.

**Conclusions:** We apply RefCell to the analysis of data from a high-throughput imaging screen of a library of 320 ubiquitin protein targeted siRNAs selected to gain insights into the mechanisms of premature aging (progeria). RefCell yields results comparable to a more complex clustering based single cell analysis method, which both reveal more potential hits than conventional average based analysis.

## Introduction

High-throughput screening (HTS) is a powerful technique routinely used in drug discovery, systematic analysis of cellular functions, and exploration of gene regulation pathways [1–4]. With modern automated microscopes, image-based HTS allows for routine imaging of thousands of cells in multiple fluorescence channels. Due to the volume and complexity of imaging data, building analysis methods has become an urgent need.

During the last decade, powerful new automated image analysis tools [5–8] that reproducibly parametrize each cell started to emerge, as well as methods for analyzing high-dimensional data specifically applicable to image-based HTS [9–19]. To identify multiple cell subtypes and quantify cellular heterogeneity, machine learning methods such as support vector machines (SVM) [15], hierarchical clustering [6], and clustering with Gaussian mixture models [9] have been introduced. While these methods are very successful in revealing cellular heterogeneity and identifying subpopulations via clustering, the “curse of dimensionality” indicates that this clustering is fraught with uncertainty: Simply as a consequence of high dimensional geometry, typical nearest neighbor distances become more and more similar to each other with increasing system dimension. Indeed, a recent study demonstrated that a number of widely used analysis approaches produce different results when applied to the same high dimensional data [20]. Furthermore, the outputs of advanced high dimensional analysis methods are not yet standardized, making comparison and interpretation of results difficult.

Here we introduce RefCell, a new method that incorporates multiple measurements simultaneously and captures similarities of cells in a single state population. RefCell is focused on the analysis of image-based HTS experiments of cellular phenotypes. Our approach captures the typical features of a single state cell population with single cell resolution. This is achieved by the introducing of “typical cells”.

We illustrate our approach in the context of an RNAi screen to identify cellular factors involved in the premature aging disease progeria. The starting point of the analysis is a set of single-cell metrics obtained through standard image-processing tools (e.g. [10, 22]). The main output of the analysis is the identification of the most significant morphological features that together provide a holistic view of the disease phenotype, and a list of significant siRNA perturbations that partially rescue the disease phenotype. We compare our pipeline to one of the more complex methods for characterizing heterogeneous cellular response [9] and found that our pipeline yields similar hits, yet is simpler, faster, and yields output graphs directly interpretable by biomedical researchers.

## Results

We demonstrate our pipeline using datasets from an image-based high-throughput siRNA screen designed to investigate cellular factors that contribute to the disease mechanism in the premature aging disorder Hutchinson-Gilford progeria syndrome (HGPS), or progeria [23], a rare, fatal disease which affects one in 4 to 8 million live births [24]. HGPS is caused by a point mutation in the *LMNA* gene encoding the nuclear structural proteins lamin A and C [25]. The HGPS mutation creates an alternative splice donor site that results in a shorter mRNA which is later translated into the progerin protein - a mutant isoform of the wild-type lamin A protein [24, 25]. HGPS is thought to be relevant to normal physiological aging as well [26–31] since low levels of the progerin protein have been found in blood vessels, skin and skin fibroblasts of normally aged individuals [29]. The progerin protein is thought to associate with the nuclear membrane and cause membrane bulging [32]. In addition to nuclear shape abnormalities and progerin expression, two additional features that have been associated with progeria are the accumulation of DNA damage inside the nucleus [33], as well as reduced and mislocalized expression of lamin B1, another lamin that functions together with lamin A [28].

These cellular hallmarks of progeria are evident at the single-cell level (Fig 2.1a; Fig S1). Typical nuclei from healthy skin fibroblasts with no progerin expression exhibit round nuclear shape, homogeneous lamin B1 expression along the nuclear boundary, and little evidence of DNA damage (Fig S1, top). In contrast, typical nuclei from HGPS patient skin fibroblasts show aberrant nuclear shape, reduced lamin B levels, and increased DNA damage (Fig S1, bottom). For a controlled RNAi screening experiment, a previously described hTERT immortalized skin fibroblast cell line was used in which GFP-progerin expression can be induced by exposure to doxycycline, causing the various defects observed in HGPS patient fibroblasts [34]. RNAi screening controls consisted of fibroblasts in which GFP-progerin expression was induced by doxycycline treatment, in the presence of 1) a non-targeting control siRNA, which allowed for full expression of GFP-progerin and formation of a progeria-like cellular phenotype in most cells, and from here on will be referred to as the GFP-progerin expressing control, or 2) a GFP-targeting siRNA, which eliminated GFP-progerin, restored a healthy-like phenotype, and from here on will be referred to as the GFP-progerin repressed control. Progerin-induced cells were plated in 384-well plates and screened against a library of 320 ubiquitin family targeted siRNAs. In addition, 12 GFP-progerin expressing controls and 12 GFP-progerin repressed controls were prepared on each imaging plate which enables estimation of control variability. Four fluorescent channels were analyzed (DAPI to visualize DNA, far-red: the nuclear architectural protein lamin B1, green: progerin, red: γH2AX as a marker of DNA damage). Images were taken at 6 different locations in each well, and each plate was imaged 4 times under the same conditions; the whole imaging procedure was applied to 4 replicate plates with identical setups (see Methods). Details of the screening process are reported in [34].

### Definition of stable classification boundaries based on typical cells

Single cell heterogeneity is prevalent in most cell population and in our screen (Fig 1). While typical progerin expressing cells exhibit reduced and inhomogeneous lamin B1 expression, pronounced DNA damage, high expression of progerin, and a blebbed cell shape, some cells in this population look like a typical healthy cell, with normal levels of homogeneously distributed lamin B1, little or no DNA damage, little to no expression of progerin, and round nuclear shape (Fig 1). Conversely, the cellular population of GFP-progerin repressed controls consists mostly of healthy-looking cells. However, a small fraction of cells in this population display features characteristic of progeria (Fig 1a). This heterogeneity is a well-established feature of HGPS patient cells [28].

**Figure 1.**
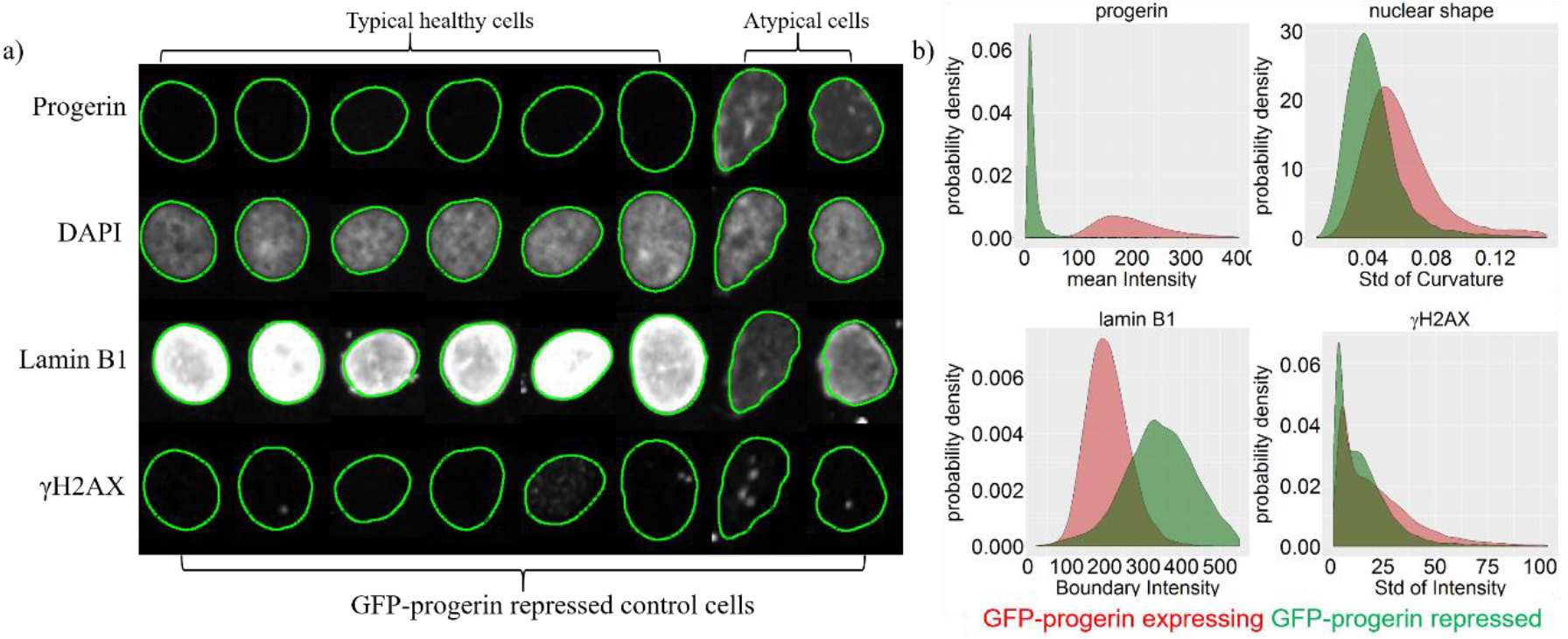
Single cell heterogeneity leads to overlapping cell populations. a) Each row corresponds to one fluorescent marker; columns show different nuclei selected from GFP-progerin repressed controls. Nuclear shapes (green contours) were extracted from the DAPI channel and mapped onto the other channels. Typical healthy cells (first six columns), exhibiting normal lamin B1 expression, little DNA damage, no expression of progerin, and round nuclear shape, as expected for GFP-progerin repressed controls. Atypical cells (two rightmost columns) exhibit characteristics of progeria, namely reduced lamin B1 expression, increased DNA damage in the γH2AX channel, expression of progerin, and blebbed nuclear shape. b) Distribution of the metric that best separates the two types of controls in each channel, based on all cells in the control samples (green: GFP-progerin repressed cells, red: GFP-progerin expressing cells). Note that the contours obtained from the DAPI channel appear slightly smaller and misaligned with the images obtained in the lamin B1 channel (see Fig S2 for the analysis of cross-channel discrepancies). The scale bar is 5 μm.

Quantification of single cell features shows the distribution of the mean intensity for all nuclei (progerin channel), the distribution of standard deviations of curvature (Lamin B1 channel), the distribution of fluorescence intensities found along the nuclear boundary (boundary intensities; Lamin B1 channel), and the standard deviation of intensities inside nucleus (YH2AX channel) (Fig 1b). These metrics were extracted via automated image analysis tools (see Methods) from all images in all control samples. For each of the four channels imaged, we show the metric that best separates GFP-progerin expressing controls (red) from GFP-progerin repressed controls (green). Except for the intensity of progerin, distributions overlap significantly, highlighting substantial heterogeneity among nuclei within each control group. The heterogeneity is largest for γH2AX, followed by nuclear shape and lamin B1.

Despite heterogeneous cellular expression, the average behavior of GFP-progerin expressing and repressed control cells are significantly different. Since the goal of this screen (as many other screens for identifying potential drugs) is to identify important perturbations that reverse the states of diseased cells to healthy-like, we focus on similarities of cells in each type of controls.

Classification of individual cells based on such overlapping distributions is challenging, as indicated by the fact that the analysis of multiple sets of 300 randomly selected cells of each of the two reference types via a Support Vector Machine (SVM) approach (see Methods) does not result in a stable classification boundary (Fig 2). To illustrate this limitation, we use 200 bootstrap samplings to identify a classification boundary using all metric dimensions simultaneously. We then extract the variability of the classification boundary in each channel (Fig 2b). We observe that classification boundaries rotated on average by more than 10 degrees between trials in the progerin channel, and by somewhat smaller amounts in the other channels.

**Figure 2.**
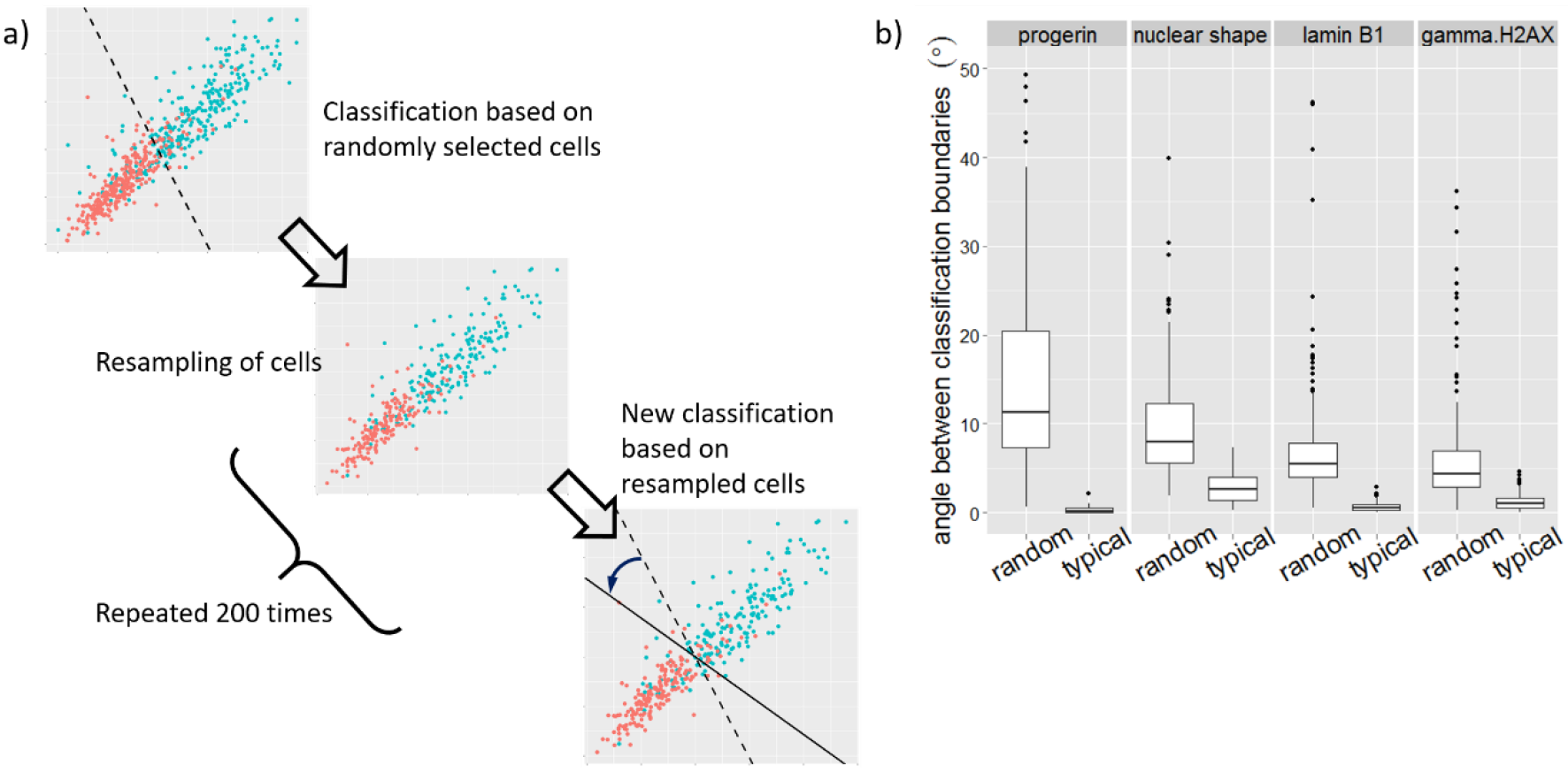
“Typical” cells yield robust metrics weighting and stable classification. a) A cartoon showing 300 randomly selected cells for each of the two control populations and a putative classification boundary. The variability in angle for 200 repeats is shown in (b). The range of angles is substantially smaller when “typical” cells are used.

Note that the angle of the classification boundary determines the relative weight of the two metrics shown in the scatter plot: for example, a vertical classification boundary indicates that the metric plotted along the vertical axis is not important for classification. Thus uncertainty about the orientation of the classification boundary implies uncertainty about the relative weight of the metrics in distinguishing both controls. To provide a reliable weighting of metrics and to find reproducible classification boundaries, we use typical cells, defined as cells close to the center of distribution of given cell population in a given channel (see Methods). Typical cells lead to stable classification boundaries with variations of less than 5 degrees in all channels (Fig 2b).

### Stable classification boundary enables identification of potential siRNA hits based on the fraction of healthy-like cells

Once a stable classification boundary is drawn based on typical healthy-like (GFP-progerin repressed control) and progeria-like (GFP-progerin expressed control) samples, all cells in all samples can be analyzed using the classification boundary. Specifically, we measured the percentage of healthy-like cells in every sample (Fig 3). We define significant siRNA perturbations, or “hits”, based on the ability of the siRNA perturbation to significantly increase the percentage of healthy-like cells (see Methods).

**Figure 3.**
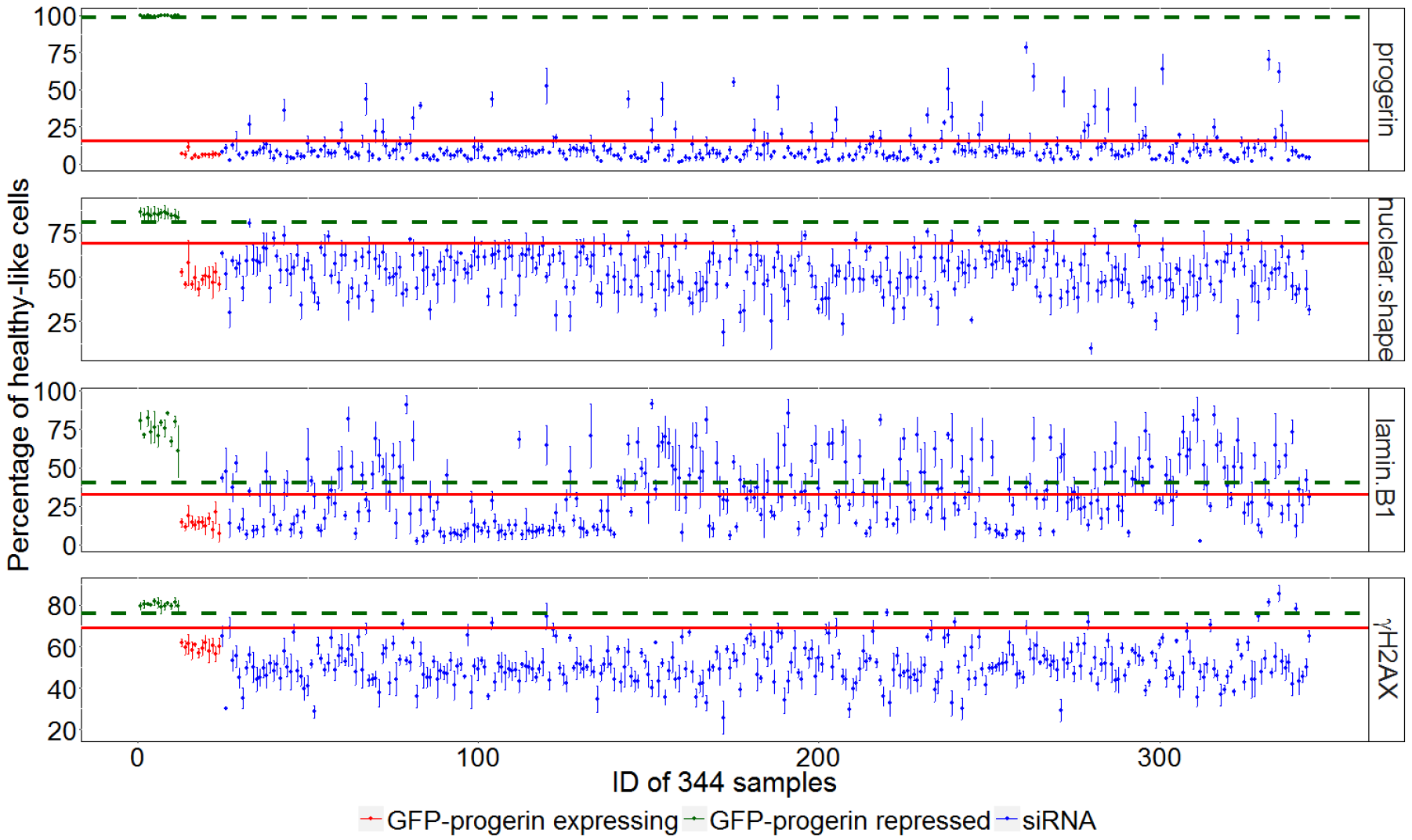
Identifying hits from the percentage of cells classified as healthy-like. A visual representation of the entire screen (320 siRNA samples, 12 GFP-progerin repressed control samples, and 12 GFP-progerin expressed control samples). Each dot represents a sample (green: GFP-progerin repressed control, red: GFP-progerin expressing control, blue: siRNA samples), with the vertical axis showing the average percentage and the error bar showing the standard deviation of healthy-like cells computed from the 4 independent replicates. False positive rate (FPR) for each siRNA is estimated from this standard deviation. The red horizontal line marks the upper boundary for GFP-progerin expressing control samples used to identify hits (5 standard deviations from the mean of all GFP-progerin expressing controls). Only siRNAs above this line with FPR < 0.05 are considered as hits. The green dashed horizontal line marks the lower boundary for GFP-progerin repressed control samples (5 standard deviations from the mean of all GFP-progerin repressed controls).

In all channels, GFP-progerin expressing and repressed controls are well separated, with the healthy-like phenotype boundary (green dashed line in Fig 3) above the hit selection threshold (red solid line in Fig 3). The separation between GPF-progerin expressing and repressed controls is the largest in the progerin channel, as expected since GFP-progerin repressed controls are derived from GFP-progerin expressing controls via GFP siRNA modulation. According to our criteria for the selection of siRNA hits (see Methods), the lamin B1 has the largest number of hits (75), followed by progerin (31), nuclear shape (8), and γH2AX (5) (see details in supplementary information S7).

The fraction of healthy-like cells in each well in the screen constitutes a metric not yet widely used in screen analysis. This metric highlights the ability of the siRNA to significantly alter some of the cells, but not all, whereas the more traditional metrics - which were also used in the original analysis of this dataset in Ref. [34] - emphasize shifts in the overall behavior. To compare the two metrics, we determine the Z-scores of the shifts in average properties (Fig 4a). Both types of Z-scores are determined based on GFP-progerin expressing control samples. For the traditional metric, the threshold is held at Z-score of 2, while our threshold is at Z-score of 5 (by Chebyshev’s inequality the probability that the hit is spurious is less than 0.04). Note that if we increase the Z-score threshold for traditional metrics to 5, there will be no hits identified. These two thresholds (gray lines) separate each panel of Figure 4a into four quadrants: perturbations identified as hits by both methods (upper right), hits identified only by traditional metrics (lower right), hits identified only by the fraction of healthy-like cells (upper left), and perturbations not identified as hits by either method (lower left). The bottom right quadrant is empty except for two siRNAs in the γH2AX channel, suggesting that our method captured nearly all hits determined by the traditional metric. On the other hand, points in the top left quadrant represent siRNA hits identified only by our approach, suggesting that our metric is more sensitive in the sense of identifying additional possible hits.

**Figure 4.**
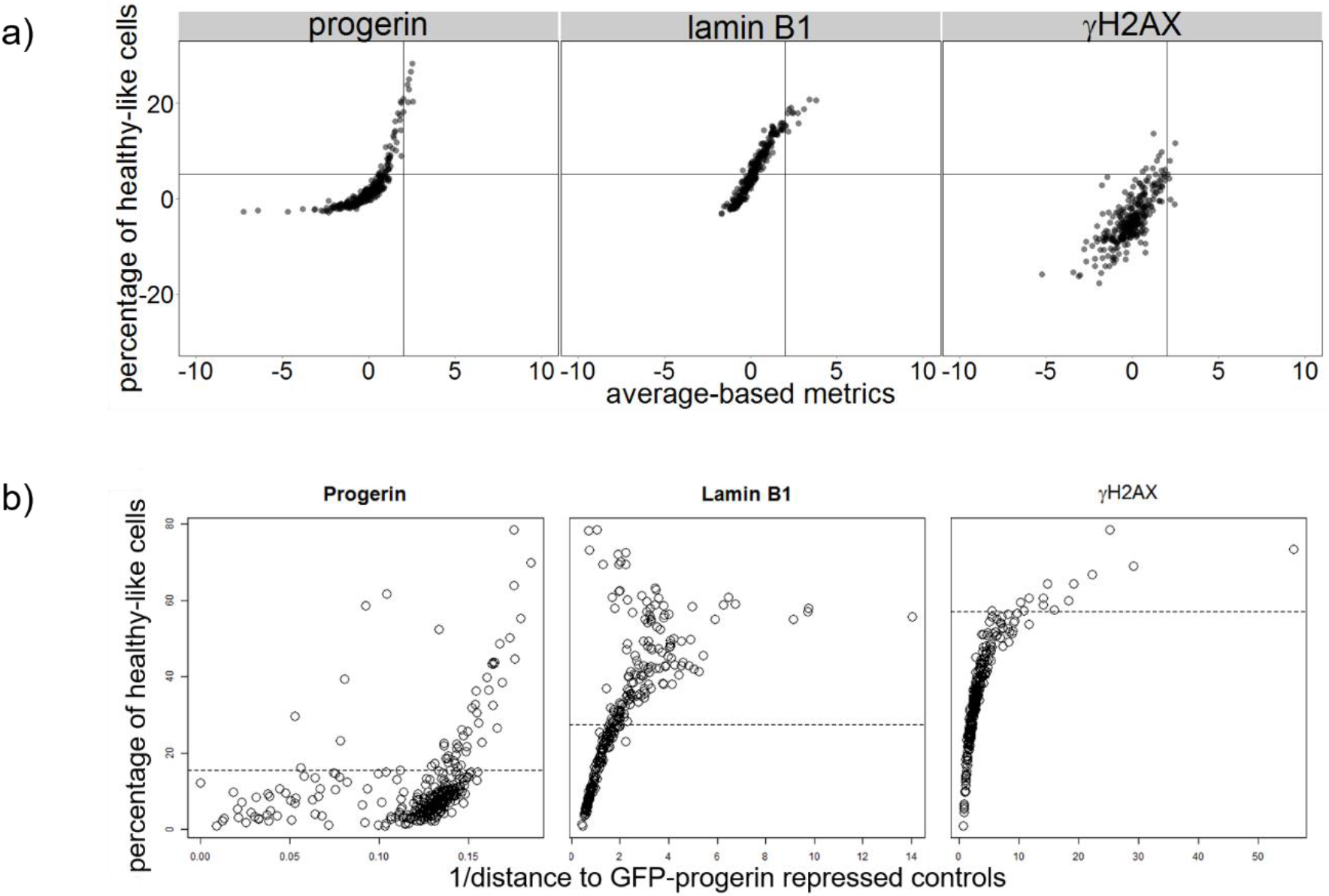
Comparing the percentage of healthy-like cells with traditional average-based metrics and another multi-dimensional analysis approach. a) Each panel depicts one channel (nuclear shape (DAPI channel) is not taken into consideration in Ref. [34], hence it is not included here). Every point represents a siRNA sample; the value shows Z-scores calculated based on the distance from the mean of all GFP-progerin expressing control samples using the traditional average-based metric calculated by directly averaging intensity measurements for all cells in a sample (x-axis) and our metric (y-axis). Gray lines indicate hit thresholds for the corresponding metrics. Note that our metric identifies every hit found by the traditional method (except for the two hits in the γH2AX channel). In addition, our metric selects additional potential hits (siRNAs in the upper left corner) missed by the traditional metric. b) Similar as in a) each panel shows one channel in the screen. Each circle depicts a siRNA sample. The horizontal axis shows inverse of the distance to GFP-progerin repressed (healthy-like) controls, the larger this value, the similar the siRNA to GFP-progerin repressed controls. The vertical axis shows the percentage of healthy-like cells, and the dashed lines are thresholds for hits in respective channels.

In addition, we benchmarked our method against one of the existing multi-dimensional analysis approaches that is also based on the difference in cell type fractions [9]. The method in Ref [9] is based on more complex clustering of all cells into multiple cell types (Fig 4b). Using the method of Ref [9], we first identified multiple clusters (9 clusters in progerin and γH2AX channels, and 8 clusters in lamin B1 channel) in 10,000 combined controls cells (5,000 for each control type). We then calculated the profile of cell distribution in each cluster for all siRNA samples and compared with GFP-progerin repressed controls (healthy-like). Since the original workflows of Ref 9 did not include hits selection, we adapted the workflow of Ref 9 and introduced the inverse distance between each siRNA sample and GFP-progerin repressed controls as the metric for hits selection. Fig 4 shows a strong correlation between the metric derived from this benchmarking test (horizontal axis) and the RefCell analysis pipeline (vertical axis) (see details in Supplementary Section 8).

### Classification boundary and metric weighting obtained via typical cells is useful for characterization of all perturbations

As explained above, we assess the phenotype for each perturbation in our high-throughput screen relative to two types of controls. Thus, the weighting of metrics given by the SVM classification boundary is based on both control phenotypes (Fig 2). In Figure 3, we had focused on subsets of cells that cross the classification boundary, i.e., that exhibit a shift in property perpendicular to the classification boundary.

In our next step, we characterize shifts of the phenotype both perpendicular and parallel to the SVM classification boundary (Fig 5a). We find that most perturbations shift cell properties perpendicular to the classification boundary. This indicates that the imaging metrics which are most important to distinguish typical cells in the two control phenotypes are also the imaging metrics that change most in the siRNA perturbations. Given that all siRNAs in this screen are ubiquitin-related (hence may affect progeria in a similar manner), this finding suggests our method really does capture the important differences between progeria phenotype and healthy phenotype. In contrast, when the classification metrics are computed from randomly selected cells - the blue points in Figure 5b - we observe shifts both parallel and perpendicular to the classification boundary. (Fig 5b). One notable exception is the progerin channel in which the two control cases are very well separated (Fig 1b).

**Figure 5.**
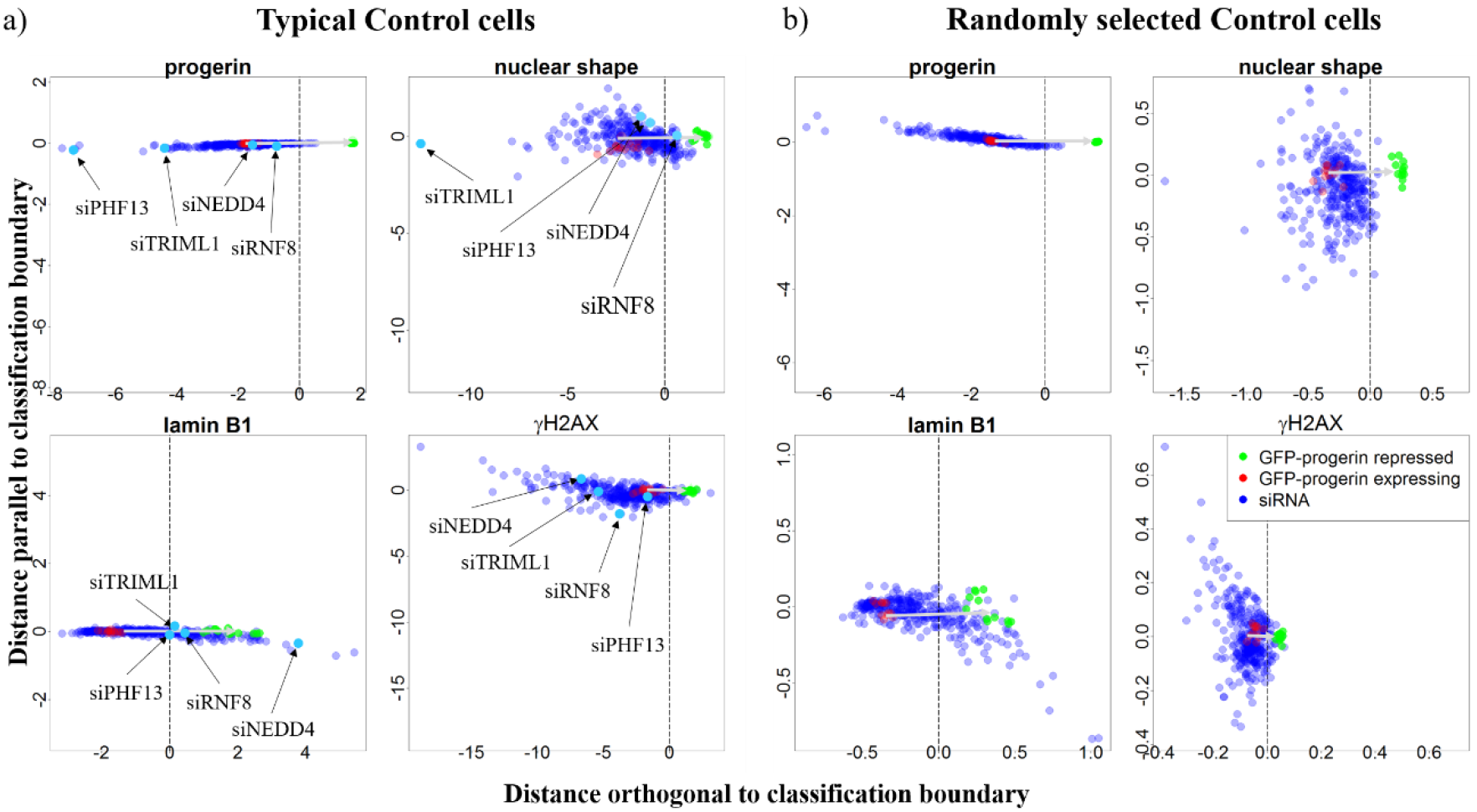
The shift of mean cell properties by siRNA perturbations for classification boundaries computed from (a) typical cells and (b) randomly selected cells. Each green and red point represents the mean of all cells in one GFP-progerin repressed (healthy-like) or GFP-progerin expressing (progeria-like) control sample, respectively. There are 12 samples for each control type. Each blue point represents the mean of all cells for one siRNA perturbation. The classification boundary is shown as a vertical dotted black line. Four siRNA samples that deviate significantly from both controls in each of the four channels are labeled (siPHF13 for progerin; siNEDD4 for lamin B1; siTRIML1 for DAPI (nuclear shape), and siRNF8 for γH2AX). Note that the range of the x-axis is the same as the range of the y-axis in all panels. **a)** Most points are preferentially shifted perpendicular to the classification boundary. Variation parallel to the classification boundary is small compared to the variation perpendicular to it. **b)** siRNA perturbations are shifted both parallel and perpendicular to the classification boundary when the classification boundary is computed from randomly selected cells.

Figure 5a also identifies siRNA perturbations that yield unusual changes in phenotype. Four examples of such siRNAs are highlighted here, one for each channel: siPHF13 for the progerin channel, siNEDD4 for the lamin B1 channel, siTRIML1 for the DAPI channel, and siRNF8 for the γH2AX channel. From each of these siRNA samples, four typical cells (picked using the same method as typical control cells; see Methods) are shown below in Figure 6 (a, b, d, and e). For comparison, four typical cells in both progeria-like and healthy-like controls are also selected (Fig 6c and f). siPHF13 treated cells (Fig 6a) express even higher levels of progerin than cells in progeria-like controls and progerin aggregates in the nucleus. Upon examining lamin B1 levels expressed by cells treated with siNEDD4 (Fig 6b), we find that lamin B1 no longer localizes only to the nuclear boundary, but spreads throughout the nucleus in an inhomogeneous way. In addition, in this case, lamin B1 expression co-localizes with progerin expression. siTRIML1 is an outlier in both the progerin and nuclear shape channel, with overexpression of progerin similar to that observed in cells treated with siPHF13. Furthermore, cells treated with siTRIML1 have nuclear shapes that are even less regular than progeria controls’. Finally, for cells treated with siRNF8 DNA damage is more substantial but also more localized (isolated bright dots in the γH2AX channel) than in progeria-like controls. These results suggest that a classification boundary built from typical cells in controls is valuable to analyze the full perturbation screen and that outliers identified in this classification point to perturbations that yield unusual properties.

**Figure 6.**
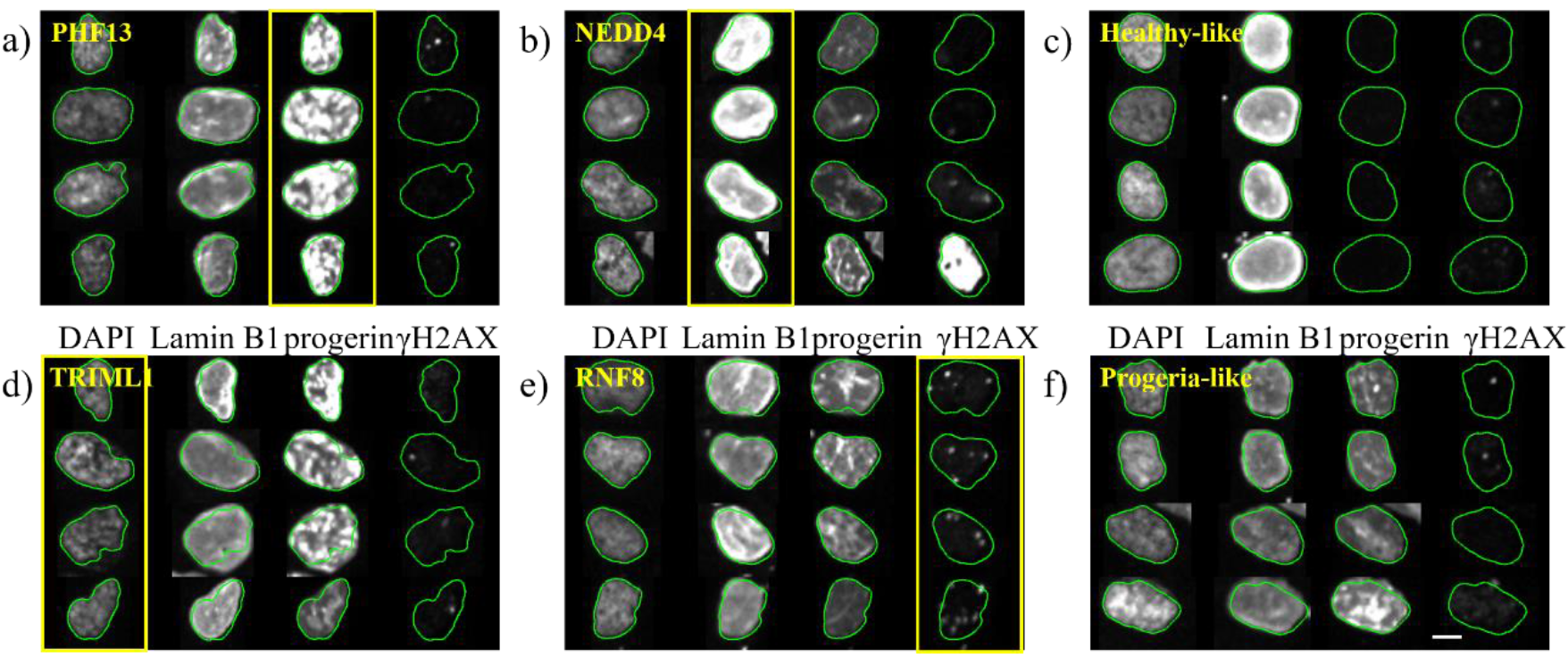
Typical cells in siRNA perturbations identified as different from both controls. a) siPHF13 is an outlier in the progerin channel: cells treated with siPHF13 express more progerin than the progeria-like control cells (f), and the expressed progerin appears to be distributed differently from the progeria control. b) siNEDD4 is an outlier in the lamin B1 channel; cells treated by siNEDD4 express more lamin B1 than the healthy-like control cells (c), and the expression is less homogenous. In addition, the expression of lamin B1 is spatially co-localized with the expression of progerin in siNEDD4-treated cells. d) siTRIML1 is an outlier in both DAPI (nuclear shape) and progerin channels. Cells treated by siTRIML1 tend to have elongated nuclei compared to the healthy-like and the progeria-like controls. Also, clusters and increased progerin expression (compared to the progeria-like control (f)) can be observed. e) siRNF8 is an outlier in the γH2AX (DNA damage) channel. Note that the contours obtained from the DAPI channel appear slightly smaller and misaligned with the images obtained in the lamin B1 channel (see Fig S2 for the analysis of cross-channel discrepancies). The scale bar is 5 μm.

### Integrating information from multiple channels increases hit detection accuracy

So far we have considered multiple metrics separately for each channel. This means that we may have labeled a cell as healthy-like based on one channel, but progeria-like when it is analyzed in another channel. This approach reflects uncertainty regarding the progeria phenotype at the single cell level: although it is known that progeria is caused by the expression of the lamin A-mutant progerin, it remains unknown how progerin expression changes other features, such as blebbed nuclear envelope, DNA damage accumulation, and mislocalized lamin B1 expression at the single-cell level, and how these different features correlate with one another. For example, in one study progeria and healthy cells were distinguished using only nuclear shape measurements [35], implying that nuclear shape is a dominant criterion in detecting progeria. However, another study found that nuclear shape could change independently from DNA damage accumulation inside the nucleus [33].

Thus, as a final analysis step, we study the relationships among the four features associated with progeria at the single-cell level. RefCell integrates single cell information from multiple channels in two different ways. First, we display the percentage of healthy-like cells for a primary marker vs. the percentage of cells identified as healthy-like according to the other three markers (Fig 7). The diameter of the circle represents the fraction of cells identified as healthy-like according to all four markers. As expected, GFP-progerin repressed controls (i.e., healthy-like controls, green circles) show a larger percentage of cells identified as healthy-like for all four markers than any of the 320 perturbations (blue circles).

**Figure 7.**
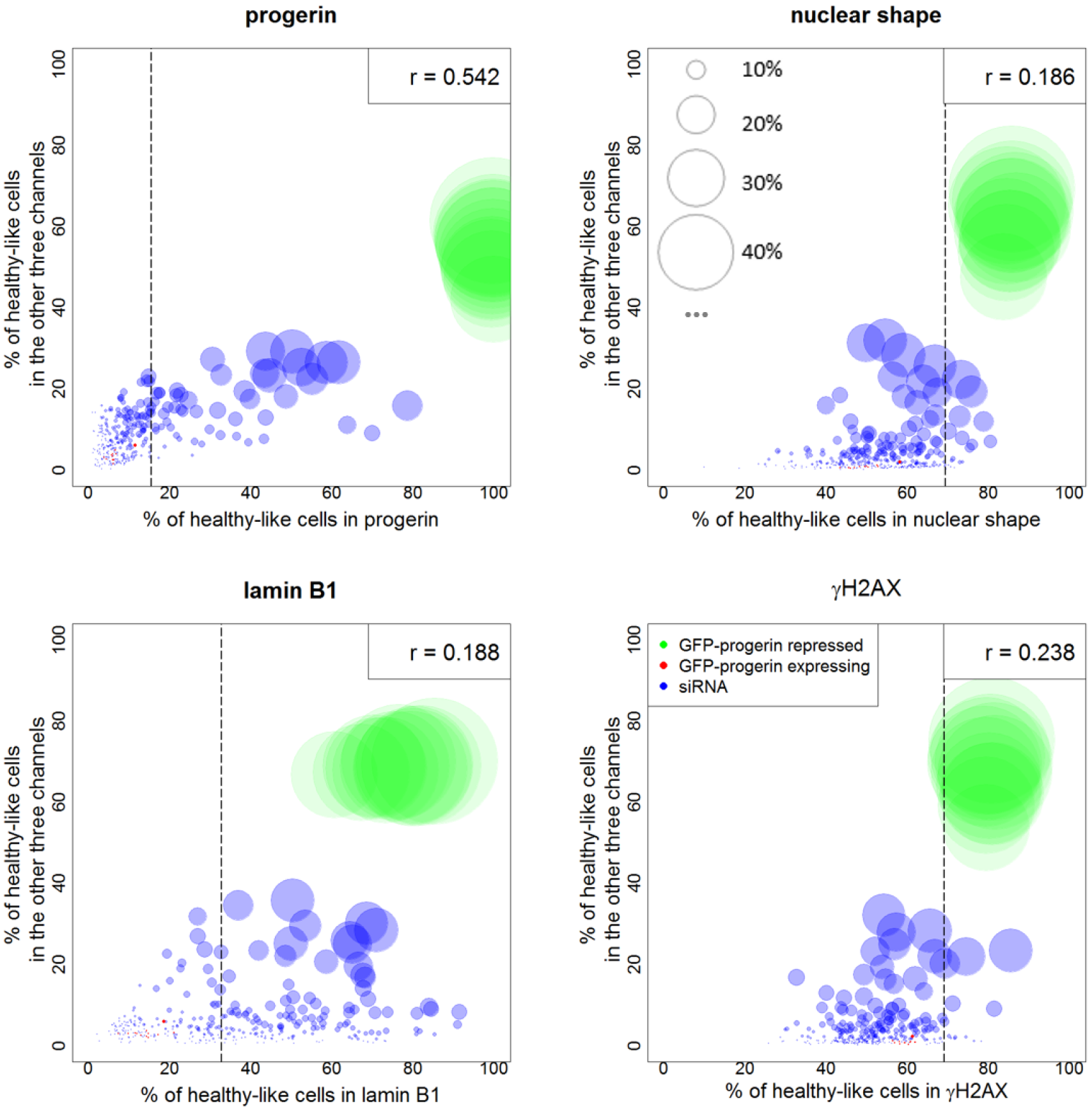
Integrating information from all channels: Percentage of healthy-like cells in one channel vs. percentage of cells classified as healthy-like in the other three channels. Each circle stands for a sample (green: GFP-progerin repressed, red: GFP-progerin expressing, blue: siRNA). The size of the circle is proportional to the percentage of cells that are classified as healthy-like in all four channels (scales are shown in top-right panel). The dashed vertical lines are thresholds for hit selection in the corresponding channel. Shown in the upper right corner of each panel is the linear correlation coefficient (note that p < 0.01 after Bonferroni correction in all cases).

Figure 7 shows that the percentage of healthy-like cells according to a given marker is correlated with the percentage identified as healthy-like according to the other three markers are correlated, although the correlation is weak in all channels except progerin.

Second, we integrated image metrics from all channels together and applied our method on combined metrics. We found that the three metrics related to progerin (mean intensity, the standard deviation of intensity and boundary intensity) are the most important metrics in separating GFP-progerin expressing and repressed controls, contributing more than 60% in the direction of classification boundary. Lamin B1 is next, contributing about 20%. In addition, we found that 99% siRNA hits identified by combining all channels are also identified by detecting hits separately for each channel; however, the combined analysis allows us to hone in on a subset of 61% of all hits (based on separate analysis of each channel).

## Discussion

One of the major usages of image-based high-throughput screening (HTS) experiments is to identify important RNAi perturbations for pathway identification or drug discovery. A major strength of image-based HTS is that measurements of multiple parameters are carried out on each cell, thus promising insights into mutual information and correlations among parameters at the single cell level. However, newly developed analysis methods yield complex and hard to interpret end results, and may actually misrepresent the data due to the “curse of dimensionality” [20]. As noted above, the “Curse of dimensionality” states that distance estimation and thus definition of nearest neighbors, which are used in clustering based high dimensional algorithms, are less meaningful in high dimensional space [36]. Here we introduce RefCell, a method that fills the gap between statistically sound average-based methods and statistically challenging high-dimensional methods. The underlying assumptions of RefCell are that the properties of typical cells are useful reference points for the biological or clinical question of interest and that the best approach to identify hits is to measure changes along a straight path (in high dimensions) between the references points.

The first step in RefCell is the selection of two sets of controls and the choice of “typical” cells within these controls. Here we choose typical cells as cells that are average in all aspects of their phenotype, i.e., all the metrics are close to the mean. In our dataset, one control represents cell nuclei of a model for progeria which show several defects, and the other control approximates healthy cell nuclei. Since image-based metrics are heterogeneous, the corresponding distributions of measured values overlap significantly at the single-cell level (Fig 1). Selecting typical cells yields distributions that are well separated, enabling stable classification boundaries between healthy-like and progeria-like cells. The classification boundary reveals both the value of each metric that marks this transition and the relative weight of each metric (Fig 2).

For the HTS used in this investigation, we find that surprisingly the metrics we identified as important are also the metrics that change most for all perturbations. A graphical representation of this observation is shown in Fig. 5a, where the two controls (green and red dots) lay out a straight path between a progeria-like phenotype and a healthy-like phenotype. All siRNA perturbations (blue dots in Fig. 5a) fall along this straight path indicating that the metrics that were identified as important are the ones that are changing most in the 320 siRNA perturbations. On the other hand, if all cells rather than typical cells are used for classification and weighting, classification boundaries are less stable (Fig. 2), and the 320 siRNA perturbations do not change the highly weighted metrics more than other metrics (the blue dots in Fig. 5b form a cloud). This indicates that the screen does not involve random perturbations but perturbations targeted specifically to progeria.

With these weights and a stable classification boundary, we were able to quantify the heterogeneity of all cells in all samples. This analysis yields a simple parameter: The fraction of cells identified as healthy-like in each sample. The fraction of normal cells had been identified in other studies as a useful parameter [37]. In RefCell, this parameter is used in multiple steps and is first determined separately for each channel to identify potential “hits” in the siRNA perturbation screen (Fig 3). RefCell then reveals a complex interplay among the four standard indicators of progeria (measured in four independent fluorescence channels), revealing that the list of hits depends strongly on the choice of indicator.

Furthermore, RefCells focus on the fraction of healthy-like cells means that any perturbation that makes a substantial fraction of cell nuclei appear healthy-like is included as a possible hit, even if the average cell properties do not change. This allows us to include all perturbations that are capable of making at least a subset of cells appear healthy-like, even if the same perturbation is ineffective or detrimental to other cells.

The final step in RefCell focuses on integrating information from multiple imaging channels (Fig 7). When considering all siRNA perturbations and all channels simultaneously, our analysis confirms that the progerin level is the most important feature in progeria disease, and that decreasing progerin expression levels is the most efficient way of removing all four principal phenotypes associated with progeria. However, we also note significant variability in how effectively a given perturbation leads to healthy-like phenotypes in each channel. This information helps prioritize hits that have been identified separately in each channel.

In addition, we compared RefCell with a published method that aims to characterize heterogeneity in cells using EM clustering with Gaussian mixture models (GMM) [9]. Since the published method did not provide a metric for hits selection, we used inverse distance to GFP-progerin repressed controls. This distance is calculated using symmetrized KL divergence as used in [9]. The higher the inversed distance, the more important the perturbation. We show that in both progerin and lamin B1 channel, our metric agrees well with the other method (see Sect S8) with Spearman correlation coefficient 0.98 for γH2AX channel and 0.91 for lamin B1 channel (p-value << 0.05 in both cases). However, the complex clustering approach does not allow us to integrate information from all channels, since complex clustering cannot be used for the analysis of cell morphology (the dimensionality of metrics is too large for meaningful clustering with “only” thousands of cell images in each sample).

In summary, RefCell represents a simple but useful computational approach for analyzing image-based HTS datasets. RefCell is broadly applicable to single-cell-based high-throughput screens that focus on perturbing cells from one distinct phenotype to another. RefCell uses image processing and machine learning algorithms to identify hits that substantially increase the fraction of cells that regain one of the two reference phenotypes. RefCell can be used to analyze each fluorescent channel separately, and also to integrate the single-cell information from all channels. Applied to a progeria HCS dataset, RefCell analysis provides robust classification boundaries between the two control groups of healthy-like and progeria-like cells, and reveals (Figure 5) that the dataset contains mostly siRNA that shift the phenotype between the two control groups. When integrating information from multiple fluorescence channels, RefCell reveals that the four standard indicators of progeria (measured in four independent fluorescence channels) are distinct, each leading to different hits in the screen.

RefCell provides a hierarchy of tools that allows step by step exploration of image-based HTS data. Starting from prioritization of metrics for each channel separately, it provides robust hits selection for each channel based on typical cells and allows for the integration of information from multiple channels. Since the key output of RefCell is visual and easy to interpret (typical cell examples, priority lists for metrics, and lists of hits), we expect that RefCell will prove valuable for a broad range of image-based high-throughput screens.

## Methods

### Experimental procedure

hTert immortalized doxycycline GFP-progerin inducible human skin fibroblasts (P1 cells) were generated and induced (96 hr). Reverse siRNA transfections were carried out in quadruplicate in a 384-well format (Perkin Elmer Cell carrier plates) in the presence of doxycycline (1 mg/ml) with pooled siRNA oligos (50nM; 4 siRNAs/target) from the Dharmacon siGENOMESMARTpool siRNA Human Ubiquitin Conjugation subset 1 and 2 libraries. Positive and negative controls consisted of GFP-targeting and non-targeting siRNA (50nM; Ambion, #AM4626, #AM4611G), respectively. Transfected cells were incubated overnight, after which 60 ml of antibiotic and doxycycline (1 mg/ml) containing medium was added, and cells were incubated for another 3 days (37 °C, 5% CO2). Details of the experiments are reported in [23]. A full list of screened siRNAs can be found in Supplementary Section 9.

### Image analysis

While metrics similar to the one used in this study could be obtained with commercial software, we used a custom image analysis method modified from methods in [38]. Details are described in Appendix S3. A list of measurements and short descriptions are shown in Table 1.

**Table 1.** Image measurements used in this study.

|  | Name of measurement | Description |
| --- | --- | --- |
| **Nuclear shape** | Area | Area of nucleus |
|  | Circularity | Ratio of perimeter to area, normalized so that a circle would have ratio 1 |
|  | Eccentricity | Eccentricity of nucleus |
|  | Invaginations | Number of invaginations along nuclear boundary |
|  | Major Axis Length | Major axis length of the best fit ellipse to nuclear boundary |
|  | Mean Curvature | Mean curvature along nuclear boundary |
|  | Mean Negative Curvature | Average of only negative curvatures along nuclear boundary |
|  | Minor Axis Length | Minor axis length of the best fit ellipse |
|  | Perimeter | Perimeter of nucleus |
|  | Solidity | Percentage of pixels inside the convex hull that are inside the boundary |
|  | Std of Curvature | Standard deviation of curvature |
|  | Tortuosity | Tortuosity of nuclear boundary |
| **Intensity** | BP Intensity | Mean intensity of points along nuclear boundary |
|  | Mean Intensity | Mean intensity inside nucleus |
|  | Std of Intensity | Standard deviation of intensity inside nucleus |

#### Selection of typical control cells

Within each control population, typical cells were defined as a core of n=300 cells closest to the mean based on the L1 (Manhattan) distance, calculated separately for each channel. We pooled all control samples together for typical control cell selection. On average there are about 20,000 cells in each type of controls. Typical progeria-like cells, selected out of the population of GFP-progerin expressed controls, show HGPS characteristic nuclear defects (increased progerin expression, misshapen nuclei, reduced lamin B1 protein levels, and increased DNA damage shown by expression of γH2AX). Typical healthy-like cells, selected from GFP-progerin repressed controls, show no sign of HGPS nuclear defects. This selection procedure was carried out independently for each replicate plate. Additional details are provided in Appendix Sect. S4.

#### Classification using Support Vector Machines (SVM)

The sets of typical cells were used to classify healthy-and progeria-like phenotypes via SVM, an efficient and robust supervised machine learning algorithm for classification [39]. Using a linear kernel, SVM finds the optimal linear boundary in instance space (straight line in 2D, planes in higher-dimensional spaces) that separates two classes of instance data points, while maximizing the margin of class separation. We performed SVM using the ksvm() function in kernlab package in R (version 3.1.1). After rescaling all nucleus metrics to zero mean and unit variance, a classification boundary was obtained between typical healthy and typical progeria cells. The distance from each nucleus to the classification boundary, which is a linear combination of all the measurements, can be used as a score to classify the proximity of that cell to each phenotype (healthy- or progeria-like). In order to distinguish between the two sides of the classification boundary, we define positive distances as associated with healthy-like cells, and negative distances with progeria-like cells. The SVM analysis also yields the relative importance of each metric in distinguishing between the two phenotypes as shown in Appendix Sect. S5.

### Identification of significant perturbations

#### Determination of the fraction of healthy-like cells

Having obtained a classifier boundary based on typical control cells, we then applied it to all samples (including all control samples and siRNA perturbations samples). For this, we first normalize all cells to be classified using the z-score transformation determined from typical control cells (i.e., subtracting the mean of typical control cells and dividing by their standard deviation). Next, we calculate the distance from each cell to the classification boundary and use the sign of the distance to classify individual cells as either healthy-or progeria-like. Finally, we calculate the percentage of healthy-like cells in each sample. This percentage is obtained separately for each replicate plate. This allows us to report the mean percentage (averaged over all replicate plates) and its estimated uncertainty (resulting from the variance over multiple replicates). The number of cells in each perturbation sample ranges from 500 to 2000. For more details, see Appendix Sect. S6.

#### Identification of siRNAs that generate significant healthy-like perturbations (“hits”)

We repeated the screen 4 times (yielding 4 independent replicates), and the analysis described above was done separately for each plate (i.e., given a sample, there are 4 independent estimates for each parameter). To carry out the hit selection process, we first averaged each parameter over the 4 replicates. Then we excluded potentially cytotoxic siRNA samples, by excluding those that contain less than 50% of cells compared to GFP-progerin repressed samples (the number of cells is similar in each sample at the start of the experiment). Next, a siRNA hit was selected based on the following two criteria: 1) the fraction of healthy-like cells is above a threshold (a mean and standard deviation were computed based on the percentage of healthy-like cells in each of the 12 GFP-progerin expressing control samples, the threshold was set to 5 standard deviations higher than the mean); 2) the false positive rate (FPR) based on the variation among the 4 replicates is less than 0.05.

## Declarations

### Availability of data and materials

The data and codes used in this manuscript are available upon request.

## References

1. Kiefer, J., et al., High-throughput siRNA screening as a method of perturbation of biological systems and identification of targeted pathways coupled with compound screening. Methods Mol Biol, 2009. 563: p. 275–87.

2. Varma, H., D.C. Lo, and B.R. Stockwell, High-Throughput and High-Content Screening for Huntington’s Disease Therapeutics, in Neurobiology of Huntington’s Disease: Applications to Drug Discovery, D.C. Lo and R.E. Hughes, Editors. 2011: Boca Raton (FL).

3. Mohr, S., C. Bakal, and N. Perrimon, Genomic screening with RNAi: results and challenges. Annu Rev Biochem, 2010. 79: p. 37–64.

4. Liberali, P., B. Snijder, and L. Pelkmans, Single-cell and multivariate approaches in genetic perturbation screens. Nat Rev Genet, 2015. 16(1): p. 18–32.

5. Inglese, J., C.E. Shamu, and R.K. Guy, Reporting data from high-throughput screening of small-molecule libraries. Nat Chem Biol, 2007. 3(8): p. 438–41.

6. Shariff, A., et al., Automated image analysis for high-content screening and analysis. J Biomol Screen, 2010. 15(7): p. 726–34.

7. Kozak, K., et al., Data mining techniques in high content screening: a survey. J Comput Sci Syst Biol, 2009. 2(04): p. 219–39.

8. Meijering, E., et al., Imagining the future of bioimage analysis. Nat Biotechnol, 2016. 34(12): p. 1250–1255.

9. Slack, M.D., et al., Characterizing heterogeneous cellular responses to perturbations. Proc Natl Acad Sci U S A, 2008. 105(49): p. 19306–11.

10. Carpenter, A.E., et al., CellProfiler: image analysis software for identifying and quantifying cell phenotypes. Genome Biol, 2006. 7(10): p. R100.

11. Jones, T.R., et al., Scoring diverse cellular morphologies in image-based screens with iterative feedback and machine learning. Proc Natl Acad Sci U S A, 2009. 106(6): p. 182631.

12. Ramo, P., et al., CellClassifier: supervised learning of cellular phenotypes. Bioinformatics, 2009. 25(22): p. 3028–30.

13. Horvath, P., et al., Machine Learning Improves the Precision and Robustness of High-Content Screens: Using Nonlinear Multiparametric Methods to Analyze Screening Results. J Biomol Screen, 2011. 16(9): p. 1059–1067.

14. Zhong, R., et al., iScreen: Image-Based High-Content RNAi Screening Analysis Tools. J Biomol Screen, 2015. 20(8): p. 998–1002.

15. Loo, L.H., L.F. Wu, and S.J. Altschuler, Image-based multivariate profiling of drug responses from single cells. Nature Methods, 2007. 4(5): p. 445–453.

16. Perlman, Z.E., et al., Multidimensional drug profiling by automated microscopy. Science, 2004. 306(5699): p. 1194–1198.

17. Jones, T.R., et al. Methods for high-content, high-throughput image-based cell screening. in Proceedings of the Workshop on Microscopic Image Analysis with Applications in Biology. 2006.

18. Birmingham, A., et al., Statistical methods for analysis of high-throughput RNA interference screens. Nature Methods, 2009. 6(8): p. 569–75.

19. Kummel, A., et al., Comparison of multivariate data analysis strategies for high-content screening. J Biomol Screen, 2011. 16(3): p. 338–47.

20. Orlova, D.Y., L.A. Herzenberg, and G. Walther, Science not art: statistically sound methods for identifying subsets in multi-dimensional flow and mass cytometry data sets. Nature Reviews Immunology, 2018. 18(1): p. 77 %@1474–1741.

21. Altschuler, S.J. and L.F. Wu, Cellular heterogeneity: do differences make a difference? Cell, 2010. 141(4): p. 559–63.

22. Verschuuren, M., et al., Accurate Detection of Dysmorphic Nuclei Using Dynamic Programming and Supervised Classification. PLoS One, 2017. 12(1).

23. Kubben, N., et al., Repression of the Antioxidant NRF2 Pathway in Premature Aging. Cell, 2016. 165(6): p. 1361–1374.

24. Capell, B.C. and F.S. Collins, Human laminopathies: nuclei gone genetically awry. Nat Rev Genet, 2006. 7(12): p. 940–52.637

25. Capell, B.C., et al., Inhibiting farnesylation of progerin prevents the characteristic nuclear blebbing of Hutchinson-Gilford progeria syndrome. Proc Natl Acad Sci U S A, 2005. 102(36): p. 12879–84.

26. Kudlow, B.A., B.K. Kennedy, and R.J. Monnat Jr, Werner and Hutchinson-Gilford progeria syndromes: mechanistic basis of human progeroid diseases. Nature Reviews Molecular Cell Biology, 2007. 8(5): p. 394.

27. Brassard, J.A., et al., Hutchinson-Gilford progeria syndrome as a model for vascular aging. Biogerontology, 2016. 17(1): p. 129–145.

28. Scaffidi, P. and T. Misteli, Reversal of the cellular phenotype in the premature aging disease Hutchinson-Gilford progeria syndrome. Molecular Biology of the Cell, 2004. 15: p. 120a–120a.

29. Zwerger, M., C.Y. Ho, and J. Lammerding, Nuclear Mechanics in Disease. Annual Review of Biomedical Engineering, Vol 13, 2011. 13: p. 397–428.

30. Allsopp, R.C., et al., Telomere Length Predicts Replicative Capacity of Human Fibroblasts. Proc Natl Acad Sci U S A, 1992. 89(21): p. 10114–10118.

31. Cao, K., et al., Progerin and telomere dysfunction collaborate to trigger cellular senescence in normal human fibroblasts. Journal of Clinical Investigation, 2011. 121(7): p. 2833–2844.

32. Goldman, R.D., et al., Accumulation of mutant lamin A causes progressive changes in nuclear architecture in Hutchinson-Gilford progeria syndrome. Proc Natl Acad Sci U S A, 2004. 101(24): p. 8963–8968.

33. Liu, Y.Y., et al., DNA damage responses in progeroid syndromes arise from defective maturation of prelamin A. Journal of Cell Science, 2006. 119(22): p. 4644–4649.

34. Kubben, N., et al., A high-content imaging-based screening pipeline for the systematic identification of anti-progeroid compounds. Methods, 2016. 96: p. 46–58.

35. Candia, J., et al., From Cellular Characteristics to Disease Diagnosis: Uncovering Phenotypes with Supercells. PLoS Comput Biol, 2013. 9(9).

36. Beyer, K.G.J.,; Ramakrishnan, R.; Shaft, U., When is “nearest neighbor” meaningful? Lecture Notes in Computer Science, 1999. 1540: p. 217–235.

37. Goransson, H., et al., Quantification of normal cell fraction and copy number neutral LOH in clinical lung cancer samples using SNP array data. PLoS One, 2009. 4(6): p. e6057.

38. Driscoll, M.K., et al., Automated image analysis of nuclear shape: what can we learn from a prematurely aged cell? Aging (Albany NY), 2012. 4(2): p. 119–32.

39. Burges, C.J.C., A tutorial on Support Vector Machines for pattern recognition. Data Mining and Knowledge Discovery, 1998. 2(2): p. 121–167.

